# Mitigating pseudoreplication and bias in resource selection functions with autocorrelation-informed weighting

**DOI:** 10.1101/2022.04.21.489059

**Authors:** Jesse M. Alston, Christen H. Fleming, Roland Kays, Jarryd P. Streicher, Colleen T. Downs, Tharmalingam Ramesh, Justin M. Calabrese

## Abstract

1. Resource selection functions are among the most commonly used statistical tools in both basic and applied animal ecology. They are typically parameterized using animal tracking data, and advances in animal tracking technology have led to increasing levels of autocorrelation between locations in such data sets. Because resource selection functions assume that data are independent and identically distributed, such autocorrelation can cause misleadingly narrow confidence intervals and biased parameter estimates.
2. Data thinning, generalized estimating equations, and step selection functions have been suggested as techniques for mitigating the statistical problems posed by autocorrelation, but these approaches have notable limitations that include statistical inefficiency, unclear or arbitrary targets for adequate levels of statistical independence, constraints in input data, and (in the case of step selection functions) scale-dependent inference. To remedy these problems, we introduce a method for likelihood weighting of animal locations to mitigate the negative consequences of autocorrelation on resource selection functions.
3. In this study, we demonstrate that this method weights each observed location in an animal’s movement track according to its level of non-independence, expanding confidence intervals and reducing bias that can arise when there are missing data in the movement track.
4. Ecologists and conservation biologists can use this method to improve the quality of inferences derived from resource selection functions. We also provide a complete, annotated analytical workflow to help new users apply our method to their own animal tracking data using the `ctmm R` package.

## 1 Introduction

Resource selection functions (RSFs) have been a mainstay of basic and applied ecology for decades (Boyce & McDonald, 1999, Manly *et al*., 2007), and are commonly used to answer questions such as “What landscape features do animals seek or avoid during their movements?” (e.g., Lamont *et al*., 2019, Prokopenko *et al*., 2017b, Valls-Fox *et al*., 2018) and “In which areas of the landscape are animals at risk of predation or disease transmission?” (e.g., Ali *et al*., 2017, Ng’weno *et al*., 2019, Rayl *et al*., 2019). They are typically conceptualized as a Poisson point process and fit in a use–availability framework, whereby environmental covariates at the locations where animals were known to be present (i.e., “used” locations) are compared with covariates at locations taken from an area assumed to be available for selection (i.e., “available” locations; Manly *et al*. 2007). Resource selection functions are usually parameterized using logistic regression, which allows researchers to easily estimate selection or avoidance of environmental features and to generate maps or data layers for use in downstream analyses (Northrup *et al*., 2013, 2021).

The relative ease of fitting RSFs has contributed to their popularity in animal ecology. However, the results of RSFs are influenced by a number of methodological challenges (Aarts *et al*., 2008, Northrup *et al*., 2013, 2021). One major (and worsening) challenge is the degree of temporal autocorrelation in tracking data, which has increased considerably as advances in tracking devices facilitate the recording of animal locations at ever-smaller intervals (Fieberg *et al*., 2010). Conventional RSFs assume that data arise from Poisson point processes and are therefore sampled independently, which means that they do not allow for any autocorrelation in the data. Autocorrelated movement data are generally less informative than independent data when estimating coarse-scale parameters, because adjacent data points share information. In other words, an animal’s location at time *t*_*i*_ is a function of both resource selection *and* its location at time *t*_*i*−1_. Estimates of resource selection are therefore overconfident and often biased when autocorrelated data are modeled as independent and identically distributed (IID). This is one form of pseudoreplication, which has long been acknowledged as a problem in ecology (Hurlbert, 1984). Modern animal tracking data are almost always positively autocorrelated (Noonan *et al*., 2019), which poses a major problem for RSFs.

Such pseudoreplication—and the incorrectly narrow confidence intervals that arise from it—can cause random variations to be incorrectly identified as significant effects. This may (at least in part) explain why a growing number of studies have documented surprising levels of apparent variation in habitat selection within species and even populations (e.g., Leclerc *et al*., 2016, Montgomery *et al*., 2018, Newediuk *et al*., 2022). Individual animals undoubtedly vary somewhat in their habitat preferences because of differences in behavioral traits (Bastille-Rousseau & Wittemyer, 2019, Leclerc *et al*., 2016, Stuber *et al*., 2022), population density (Avgar *et al*., 2020, Matthiopoulos *et al*., 2015, van Beest *et al*., 2014), and habitat availability (Aarts *et al*., 2013, Godvik *et al*., 2009, Mysterud & Ims, 1998). However, substantial levels of Type II error in selection parameters can easily arise when pseudoreplication from autocorrelated data causes unimportant covariates to be misleadingly found to be significant. Accurate quantification of uncertainty is therefore paramount for accurate inference concerning habitat selection. In addition, RSFs that assume IID data are unable to account for varying levels of autocorrelation in movement data. This might happen when environmental covariates inhibit successful estimation of animal locations (e.g., dense vegetation blocking Global Positioning System [GPS] satellite reception), leaving unsampled gaps in an animal movement track (Fleming *et al*., 2018, Frair *et al*., 2010, Lewis *et al*., 2007). When the gaps in data are correlated with specific covariates of interest, the level of autocorrelation in animal locations is also correlated with those covariates. Despite widespread awareness of these problems, no generally accepted solutions have been described, and researchers often fit IID RSF models to autocorrelated tracking data.

Historically, data thinning has been the most common suggestion for reducing autocorrelation in tracking data used to inform RSFs (e.g., Hooten *et al*., 2014, Northrup *et al*., 2013, Swihart & Slade, 1985). While pragmatic and easy to implement, data thinning suffers from a number of limitations as a method of mitigating autocorrelation. First, thinning inherently requires discarding data, which results in the loss of useful information and can lead to imprecise parameter estimates. Second, thinning typically involves using statistical tests for the presence of statistically significant levels of autocorrelation—a target which is both arbitrary and not optimized for the task of estimating distributions (Aarts *et al*., 2008). Under-thinning results in misleadingly narrow confidence intervals and possibly biased coefficient estimates, while over-thinning results in an unnecessary loss of information and statistical efficiency. Third, highly irregular time-series, such as those obtained from tracking aquatic animals (Breed *et al*., 2011, Fleming *et al*., 2018) or sampling designs targeted at identifying behaviors at multiple temporal scales (e.g., Scantlebury *et al*., 2014, Ullmann *et al*., 2020), may not be particularly amenable to thinning.

Increasingly, researchers use step-selection functions (hereafter SSFs; Avgar *et al*., 2016, Fortin *et al*., 2005, Thurfjell *et al*., 2014) to account for autocorrelation in animal movements during studies of resource selection (e.g., Alston *et al*., 2020, Dickie *et al*., 2020, Kohl *et al*., 2018, Merkle *et al*., 2016, Prokopenko *et al*., 2017a). This approach builds upon conventional RSFs by generating available points (or movement steps) using a biologically realistic, movement-informed sampling scheme. However, SSFs still have important limitations. First, because available points in conventional SSFs depend on distributions of step lengths and turning angles on a fixed sampling interval, conventional SSFs cannot handle irregularly sampled data—SSFs can only be parameterized using animal location data that were sampled at a constant sampling interval (but see Brost *et al*. (2015), Christ *et al*. (2008), and Johnson *et al*. (2008) for examples of seldom-used but conceptually similar models that can accommodate missing locations). Second, parameters of SSFs are explicitly scale-dependent—parameters change as the sampling schedule of a movement track changes (Avgar *et al*., 2016, Fieberg *et al*., 2021, Signer *et al*., 2017). In other words, if the same movement track were to be sampled at different intervals, the SSFs parameterized using the different samples would provide different estimates of selection. Third, SSFs do not invariably mitigate the pseudoreplication that arises from treating autocorrelated data as IID—step selection functions assume that steps are independent, so when the sampling interval is smaller than the length of time required for steps to be statistically independent of one another, SSFs will also produce misleadingly narrow confidence intervals. Additional measures such as data thinning (Babin *et al*., 2011, Robb *et al*., 2022), variance inflation (Nielsen *et al*., 2002), generalized estimating equations (Craiu *et al*., 2008, Prima *et al*., 2017), and modeling interactions between steps can be employed to widen confidence intervals associated with SSF parameters, but these methods lack an objective and easily implementable target for the appropriate width of confidence intervals.

Likelihood weighting offers another potential solution for mitigating the adverse effects of temporal autocorrelation on RSFs. In this framework, the contributions of individual animal locations are weighted according to a joint likelihood function, such that the sum of all weights matches an “effective sample size” derived from stochastic process models that describe the autocorrelation structure of an animal movement track. This framework avoids two problems caused by data thinning. First, instead of considering one thinned subsample of data and its set of resulting parameter estimates, or several thinned subsamples of the data and an average over their parameter estimates (which would reduce noise—and potentially bias—arising from thinning), all subsamples of independent data and an average of their log-likelihoods are evaluated during parameter estimation. In other words, autocorrelated data are down-weighted rather than discarded. Second, instead of thinning the data to an *ad hoc* threshold (e.g., less than 5% autocorrelation), data are weighted using an objective estimate of “effective sample size” that quantifies confidence in a home-range estimate. As an additional benefit, likelihood weights can also account for sampling biases that arise from irregular sampling. Such bias can occur, for example, when covariates are over- or under-sampled, such as when environmental covariates are associated with differential probability of successful triangulation of animal locations (Fleming *et al*., 2018, Frair *et al*., 2010, Lewis *et al*., 2007). This means that irregular sampling of non-independent data (which is incompatible with the IID assumption underlying conventional RSFs) can be explicitly accounted for in an RSF model.

In this study, we introduce a method for autocorrelation-informed likelihood weighting of animal locations to mitigate pseudoreplication and bias in parameter estimates of RSFs. We provide a description of the mathematical principles underlying our method, demonstrate its practical advantages over conventional approaches using simulations and empirical animal tracking data for a water mongoose (*Atilax paludinosus*), a caracal (*Caracal caracal*), and a serval (*Leptailurus serval*), and discuss pathways for continuing to improve upon our method. We also provide an annotated analytical workflow for using the `ctmm R` package (Calabrese *et al*., 2016) to apply our method to animal tracking data (Appendix 1).

## 2 Materials and Methods

### 2.1 Mathematical Concepts and Definitions

In a conventional RSF, the log-likelihood of *n* IID samples of an inhomogeneous Poisson point process model is given by

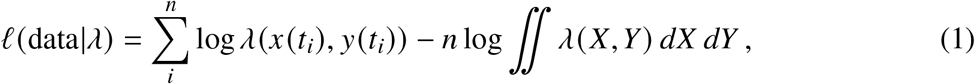

where *λ* (*x, y*) is the intensity function, which contains all model parameters, and where, for simplicity, we have assumed that *λ* does not change in time. This likelihood function is typically approximated numerically via weighted logistic regression, and *λ* is usually constructed to be an exponential model (Fieberg *et al*., 2021, Northrup *et al*., 2021), but other approaches have also been proposed and used (e.g., Cooper & Millspaugh, 1999, Lele & Keim, 2006, Nielson & Sawyer, 2013).

Within *λ*, we include area terms (*x, y*, and *x*^2^ + *y*^2^) that produce a bivariate Gaussian distribution of animal locations, so that when no resource selection occurs, the weighted likelihood reproduces a Gaussian home range estimate that approximates the maximum likelihood estimate of the autocorrelation model. This Gaussian distribution becomes the null RSF model (i.e., the domain of availability, and the area of movement when there is no habitat selection; Fieberg *et al*. 2021, Horne *et al*. 2008). The effective sample size is roughly the number of times an animal has crossed the linear extent of its home range, which dictates how well the area and shape of the Gaussian home range estimate can be resolved from the sampled movement path of an animal (Fleming *et al*., 2019). When selection parameters are supported, then we obtain a synoptic model of habitat selection (*sensu* Horne *et al*., 2008) within a Gaussian area. Further details of this configuration of the RSF model can be found in Appendix 3.

To the conventional IID log-likelihood, we incorporate our RSF log-likelihood weights, which are given by

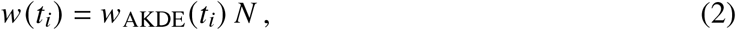

where *w*_AKDE_(*t*_*i*_) denotes weights optimized for non-parametric autocorrelated kernel density estimation at each sampled location (Fleming *et al*., 2018), which sum to 1. *N* ≤ *n* is the effective sample size of the autocorrelated Gaussian area estimate, so that our weights, *w* (*t*_*i*_), sum to *N*. The AKDE weights minimize error in non-parametric kernel density estimation, where for a sample of *n, q*-dimensional locations **r**(*t*_*i*_) at times *t*_*i*_, a weighted kernel density estimate can be represented as

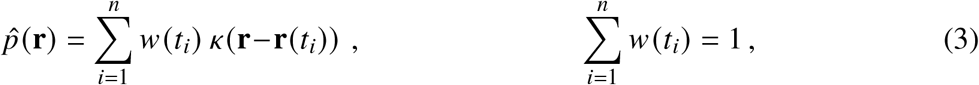

where *κ* denotes a Gaussian kernel with covariance ***σ***_B_ (the bandwidth matrix). Both the bandwidth matrix and the weight vector **w** with *w*_*i*_ = *w* (*t*_*i*_) are optimized to minimize the mean integrated square error

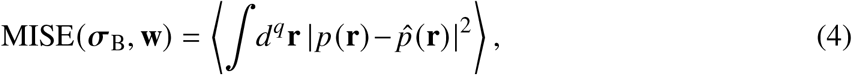

where ∫*d*^*q*^**r** denotes the *q*-dimensional volume integral and (· · ·) denotes the expectation value with respect to the distribution of the data, **r**(*t*_*i*_), which may be autocorrelated. Because the true density function *p* (**r**) is unknown and kernel density estimation is non-parametric, the MISE (4) must be approximated, and different approximations correspond to different methods of kernel density estimation—all being asymptotically optimal (Izenman, 1991, Silverman, 1986, Turlach, 1993).

The inclusion of area terms in the RSF model is a form of parametric Gaussian density estimation, which allows weights to serve a similar purpose in RSFs as for kernel density estimation. Weights (2) also provide identical estimates of the mean location in both kernel and Gaussian density estimation, indicating that they provide a bias correction for the Gaussian model. The weights *w* (*t*_*i*_) are used to weight each sampled location so that the final log-likelihood is of the form

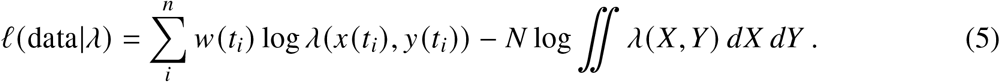

These weights can be used to re-weight data points for any method of approximating inhomogeneous Poisson point processes, but details of implementation will vary for different methods and software platforms. Likelihood (5) can be viewed as an example of a ‘composite likelihood’ (Varin *et al*., 2011) that produces an unbiased estimating equation for the mean location, but with the additional property that the total weight is fixed to the effective sample size.

### 2.2 Simulations

To validate our theoretical argument and demonstrate the value of our approach on animal location data, we performed three sets of simulations. For all three sets of simulations, we started by simulating movement paths for animals following an isotropic Ornstein-Uhlenbeck Foraging movement model (Fleming *et al*., 2014) with a location variance (*σ*) of 200,000 m^2^, velocity persistence timescale (*τ*_*v*_) of 1/3 day, and home range crossing timescale (*τ*_*p*_) of 1 day. This is equivalent to an animal with a circular home range of 3.76 km^2^, with three correlated bouts of movement each day, and which crosses its home range roughly once per day. Because range crossings occurred daily, the effective sample size in this movement track is equivalent to the number of days the movement track is sampled, and sampling locations from the movement track more frequently than once per day yields autocorrelated data. We overlaid each movement track on a raster consisting of equal amounts of two habitat types (hereafter, Habitat 1 and Habitat 2) that each take up half of the raster surface. All simulations were performed using the `ctmm` R package (v0.6.2; Calabrese *et al*., 2016) in the R statistical software environment (v3.6.2; R Core Team, 2020). Habitat rasters were created using the `raster` R package (v3.4-10; Hijmans, 2021).

#### Scenario λ: Absolute vs. Effective Sample Size

We first simulated a scenario that demonstrates that our weighting scheme does not reduce bias simply by inflating confidence intervals such that it becomes impossible to identify habitat selection when it occurs. For this scenario, we simulated habitat selection occurring by arraying habitat in 10 m vertical stripes and then simulated habitat-independent movement tracks over this layer. We sampled these tracks while varying (1) the sampling rate (2, 4, 8, 16, and 32 times per day for 90 days) or (2) the sampling duration (12 times per day for 16, 32, 64, 128, and 256 days). We then shifted every other location in Habitat 1 further right by 10 m, which moved those locations into Habitat 2. This alteration changed the fit of the underlying movement model very little, but led to the animals being located in Habitat 2 roughly three times as often as they were located in Habitat 1, emulating habitat selection. Finally, we used weighted RSFs to estimate strength of selection for Habitat 2 compared with Habitat 1, and the confidence intervals around this parameter estimate. In this scenario, parameter estimates should be constant around the expected parameter (*ln*(3)) in all scenarios, but confidence intervals should only meaningfully contract when the effective sample size (i.e., sampling duration) increases, and not just when the temporal resolution of the data increases.

#### Scenario 2: Data thinning

We then simulated a scenario that demonstrates that data thinning creates substantially more variation around RSF parameter estimates than weighting. As in Scenario 1, we simulated habitat selection occurring by arraying habitat in 10 m vertical stripes and then simulated a habitat-independent movement track over this layer. The track was sampled 50 times per day for 64 days. We then shifted every other location in Habitat 1 further right by 10 m, which moved those locations into Habitat 2. We then created 50 subsampled movement tracks (all possible regular subsamples of this data in which the absolute sample size is roughly equal to the effective sample size, rendering locations independent from one another). We used a weighted RSF to estimate strength of selection for Habitat 2 compared with Habitat 1, and the confidence intervals around this parameter estimate, for the full movement track. We used IID RSFs to estimate strength of selection for Habitat 2 compared with Habitat 1, and the confidence intervals around this parameter estimate, for each subsampled movement track. In this scenario, the parameter estimates for the weighted RSF should be near the expected parameter (*ln*(3)), but thinning the data will create substantial variation in IID parameter estimates because of loss of information about the movement track.

#### Scenario 3: Triangulation Failure

We finally simulated a scenario in which bias could arise when certain habitat covariates of interest cause disproportionate amounts of triangulation failure in GPS devices. This is known to happen, for example, in rugged terrain, thick vegetation, and underwater (e.g., Breed *et al*., 2011, O’Neill *et al*., 2020, Streicher *et al*., 2021). For this scenario, we simulated no habitat selection occurring (i.e., the animal movement path was simulated independently of the habitat raster), but we censored every other location in Habitat 2, which would be consistent with a habitat-specific triangulation success rate of 50% (an extreme case of habitat-specific triangulation failure). Because the spatial configuration of habitat affects estimates of resource selection (Northrup *et al*., 2013), we configured our habitat rasters in two different ways to illustrate how weighting responds to habitat configuration: (1) a highly clustered case, in which the two habitats were arrayed in two large contiguous blocks, and (2) an unclustered case where the habitats were arrayed in alternating vertical stripes that were 10 m in width (identical to the raster in Scenarios 1 and 2). We sampled these tracks 1, 2, 4, 8, and 16 times per day, for 90 days, on 400 movement tracks in each habitat configuration. We finally used weighted and IID RSFs to estimate strength of selection for Habitat 1 compared with Habitat 2, and the confidence intervals around this parameter estimate.

This scenario demonstrates two useful properties of our weighting scheme. First, by varying the sampling rate, we can demonstrate that an RSF that treats data as IID becomes increasingly confident in its parameter estimates as data become more autocorrelated—even when they are biased—while weighted RSFs maintain stable confidence intervals as locations are sampled more frequently because the effective sample size of the data set is not increasing. In other words, reducing the time interval between sampled animal locations causes the confidence intervals around selection parameters to become narrower in IID RSFs (because increasing autocorrelation in the data causes pseudoreplication) but not in weighted RSFs (because autocorrelation is explicitly modeled and accounted for). Second, by simulating habitat-specific location failure, we demonstrate that this weighting scheme can reduce sampling-induced bias in RSF parameter estimates compared with RSFs that assume data are IID. An IID RSF should detect a pattern of selection of Habitat 1 over Habitat 2 that is a statistical artefact of triangulation failure; a weighted RSF will shift the parameter estimate toward zero and expand the confidence intervals around the parameter estimate, correctly inferring that no habitat selection is occurring when effective sample sizes are large enough to reliably estimate parameters. However, its ability to do this depends on the degree of spatial clustering of habitat covariates—which is known to influence RSF parameter estimates (Northrup *et al*., 2013, Street *et al*., 2021). As more consecutive locations occur within a single habitat type, the average weights of locations in the habitat type with frequent triangulation failure become larger, allowing the model to reduce the influence of the habitat-specific sampling intervals on parameter estimates.

### 2.3 Empirical Examples

To demonstrate the application of our method on real-world animal location data, we performed IID and weighted RSFs on tracking data from a water mongoose (144 locations over 58 days; Fig. S1), a caracal (504 locations over 85 days; Fig. S2), and a serval (3603 locations over 321 days; Fig. S3). All animals were tracked using GPS telemetry; further details on the specifics of animal capture and tracking can be found in Streicher *et al*. (2021), Ramesh *et al*. (2016a), and Ramesh *et al*. (2016b). All three animals are missing a meaningful number of locations because of GPS triangulation failure. For each animal, we estimated the effects of two land cover types (built-up [human settlements, roads, railways, and airfields] and plantation [exotic tree plantations]) on their habitat selection compared with a grouped reference category consisting of the remaining land cover types. For our land cover data, we used a land-use map with 20-m resolution from Ezemvelo KZN Wildlife (EKZN Wildlife & GeoTerraImage, 2018) reclassified into the three categories of interest (i.e., plantation, built-up, and reference).

For each animal, we used variogram analysis (Fleming *et al*., 2014) to ensure animals were range-resident, fit and selected an autocorrelated movement model that best described the animal’s movements using perturbative Hybrid Residual Maximum Likelihood (phREML; Fleming *et al*., 2019) and Akaike’s Information Criterion corrected for small sample sizes (AICc), estimated utilization distributions for each animal (wAKDE for weighted RSFs [Fleming *et al*. 2018]; conventional KDE for IID RSFs [Worton 1989]), and fit RSFs using the `ctmm` R package (v0.6.2; Calabrese *et al*., 2016) in the R statistical software environment (v3.6.2; R Core Team, 2020).

For the caracal, we then examined how using IID RSFs on thinned data sets would alter inferences gained from this data set. To do this, we created 27 subsampled movement tracks (all possible regular subsamples of this data in which the absolute sample size is roughly equal to the effective sample size, rendering locations independent from one another). We used a weighted RSF to estimate strength of selection for built-up areas (accounting for selection for plantation), and the confidence intervals around this parameter estimate, for the full movement track. We used IID RSFs to estimate strength of selection for built-up areas (accounting for plantation), and the confidence intervals around this parameter estimate, for each subsampled movement track.

## 3 Results

### 3.1 Simulations

#### Scenario 1: Absolute vs. Effective Sample Size

In our simulations of habitat selection, we found that weighted RSFs performed as expected, with parameter estimates of weighted RSFs remaining consistently around the expected parameter (*ln*(3); Fig. 1). The width of confidence intervals remained stable as the sampling rate (i.e., autocorrelation) increased (Fig. 1A), but contracted as sampling duration (i.e., effective sample size) increased (Fig. 1B). Weighted RSFs could detect habitat selection when effective sample sizes were ≥ 32 (or 32 days) in this example.

**Figure 1:**
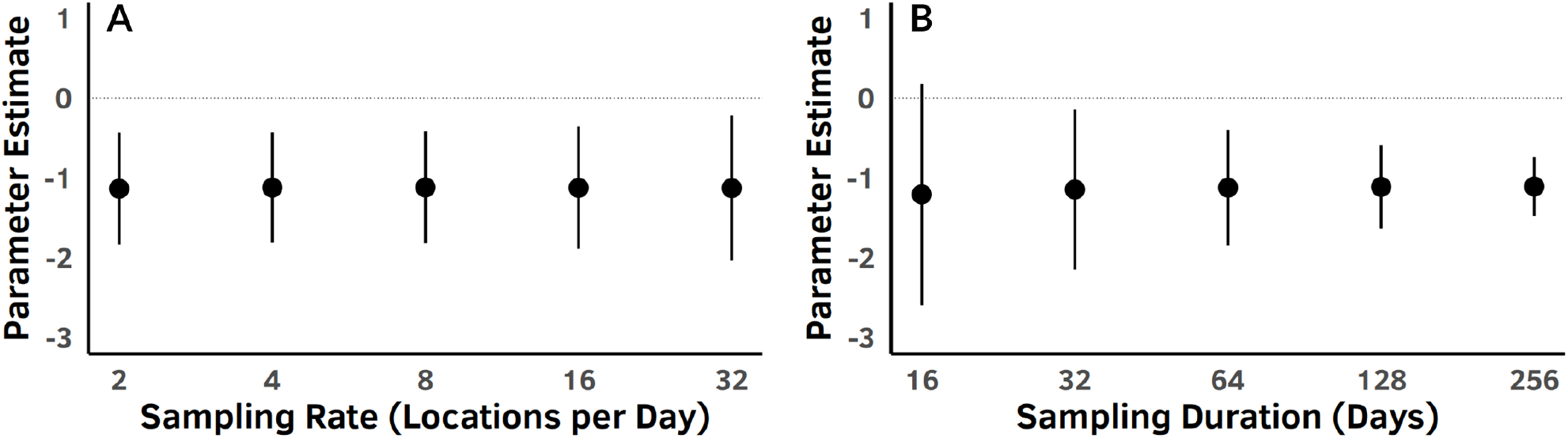
Mean parameter estimates and confidence intervals around those parameter estimates for weighted RSFs during simulations of habitat selection. In this case, the animal is selecting habitat such that its movement path contains three times as many locations in Habitat 2 as in Habitat 1. Panel A shows the performance of weighted RSFs as the sampling rate (i.e., autocorrelation) increases, while Panel B shows the performance of weighted RSFs as sampling duration (effective sample size) increases. As expected, point estimates of the selection parameter are constant across all scenarios, while confidence in those parameters increases as sampling duration (i.e., effective sample size), but not sampling rate (i.e., autocorrelation), increases.

#### Scenario 2: Data Thinning

In our simulations of data thinning, we found that RSFs performed as expected, with the parameter estimates of the weighted RSF very close to the expected parameter (*ln*(3)) while the IID RSFs paramaterized using thinned data sets exhibited substantial variation in parameter estimates (range: [-1.94, -0.44]; Fig. 2). The expected parameter fell within 48 out of 50 (96%) of the IID RSF 95% confidence intervals, indicating that our measure of effective sample size is a reasonable measure of independence. One IID RSF failed to identify statistically significant habitat selection.

**Figure 2:**
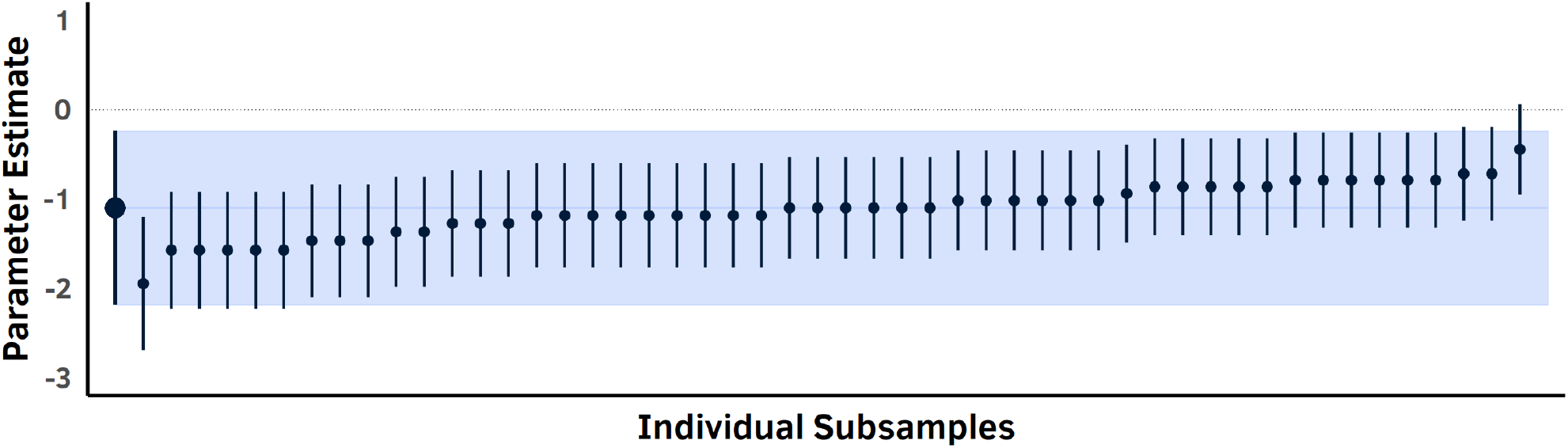
Parameter estimates and confidence intervals around those parameter estimates for one weighted RSF performed on a complete movement track and 50 IID RSFs performed on subsamples of the movement track in which each sequential location was independent. In this case, the animal is selecting habitat such that its movement path contains three times as many locations in Habitat 2 as in Habitat 1. The larger point and bar on the far left represent the weighted RSF, while the smaller points and bars represent individual subsets. The light blue box outlines the confidence interval of the weighted RSF, while the darker blue horizontal line identifies the expected parameter (ln(3)). As expected, IID RSFs parameterized using thinned data sets exhibit substantial variation in their estimated parameters.

#### Scenario 3: Triangulation Failure

In our simulations of triangulation failure, we found that weighted and IID RSFs performed as expected. Across 400 individual movement paths, parameter estimates of weighted RSFs were closer to zero than parameter estimates of IID RSFs (Fig. 3; Table 1). Confidence intervals around parameter estimates of weighted RSFs contained zero much more often (Table 1, top half; Figs. 3A,C) than confidence intervals around parameter estimates of IID RSFs (Table 1, bottom half; Figs. 3B,D). Confidence intervals around parameter estimates of IID RSFs became narrower as the sampling rate (and thus autocorrelation) increased, while the width of confidence intervals around parameter estimates of weighted RSFs remained similar across all sampling rates (Fig. 3). Notably, weighted RSFs were better able to successfully identify a lack of habitat selection (i.e., weighted RSF parameter estimates became increasingly closer to zero and a higher proportion of confidence intervals contained zero) as the sampling rate increased, while IID RSFs performed worse as the sampling rate increased (i.e., parameter estimates were largely consistent, but confidence intervals around the [biased] estimates contracted sharply as the sampling rate increased). The benefits of weighted RSFs are also enhanced when habitat is more highly clustered, but likelihood weighting still improves both parameter estimates and confidence intervals when habitat is unclustered (Table 1).

**Table 1:** The effect of sampling rate and habitat clustering on parameter estimates, the width of confidence intervals, and percentage of non-significant parameter estimates of weighted and IID RSFs across 400 simulated movement paths. Movement paths occurred with no habitat selection but with habitat-specific triangulation failure. Parameter estimates of weighted RSFs are closer to zero than for IID RSFs, confidence intervals around wRSF parameter estimates are wider and more consistent than for IID RSFs, and wRSFs are more likely to identify that there is no habitat selection occurring when location data are autocorrelated.

| RSF Type | Habitat Clustering | Sampling Rate | Parameter Est. | % Non-significant |
| --- | --- | --- | --- | --- |
| wRSF | High | 1 | -0.59 | 83.8 |
|  |  | 2 | -0.43 | 93.5 |
|  |  | 4 | -0.23 | 99.3 |
|  |  | 8 | -0.10 | 99.3 |
|  |  | 16 | -0.07 | 99.5 |
|  | Low | 1 | -0.60 | 51.5 |
|  |  | 2 | -0.55 | 46.3 |
|  |  | 4 | -0.50 | 58.0 |
|  |  | 8 | -0.49 | 60.8 |
|  |  | 16 | -0.49 | 62.5 |
| IID | High | 1 | -0.68 | 61.0 |
|  |  | 2 | -0.68 | 37.5 |
|  |  | 4 | -0.71 | 15.8 |
|  |  | 8 | -0.71 | 7.3 |
|  |  | 16 | -0.74 | 3.8 |
|  | Low | 1 | -0.69 | 24.3 |
|  |  | 2 | -0.68 | 1.5 |
|  |  | 4 | -0.69 | 0.0 |
|  |  | 8 | -0.69 | 0.0 |
|  |  | 16 | -0.69 | 0.0 |

**Figure 3:**
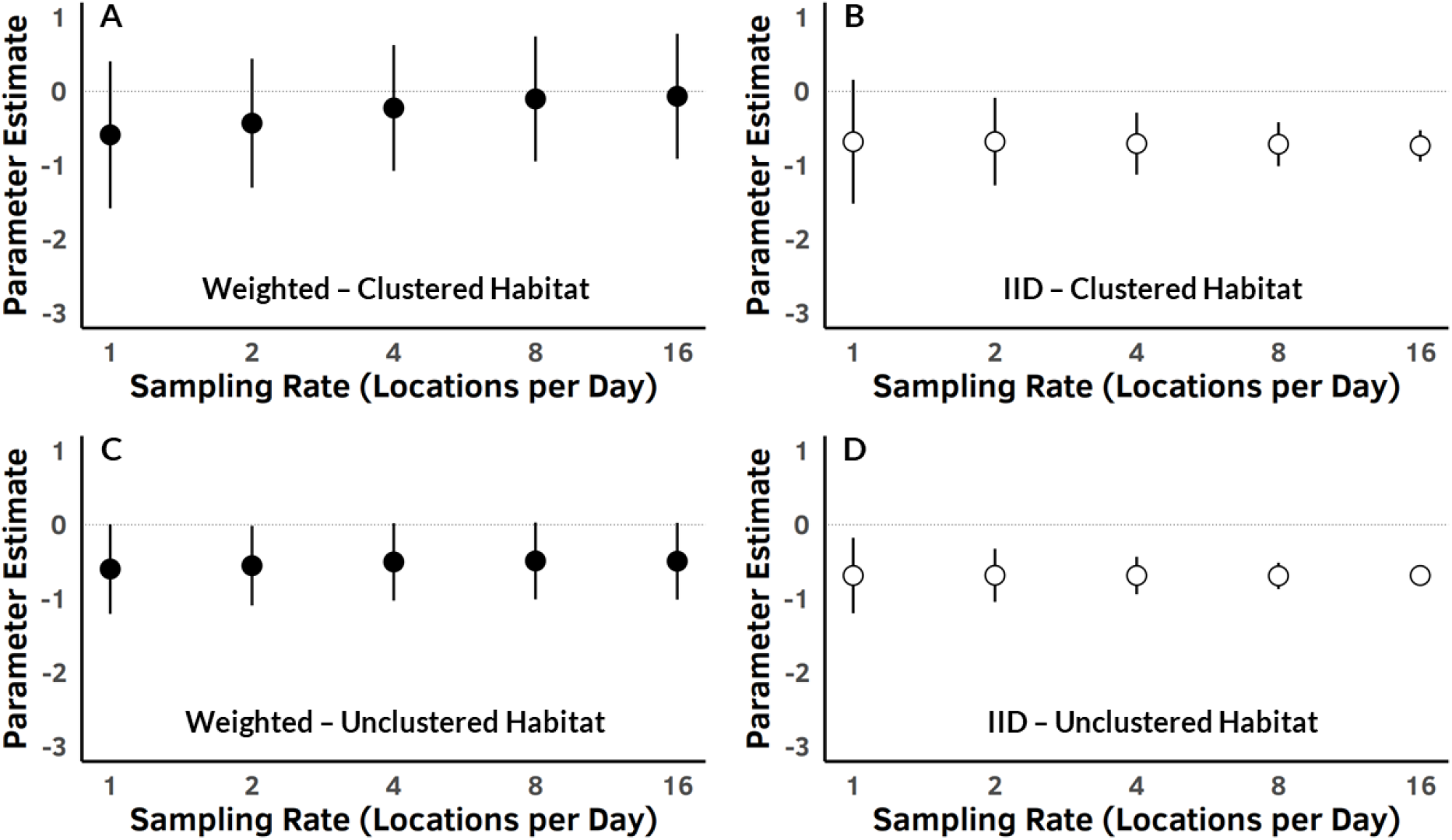
Mean parameter estimates and confidence intervals for weighted RSFs vs. IID RSFs for 400 simulations of habitat-specific GPS triangulation failure. Row 1 (Panels A and B) shows the performance of RSFs in a scenario of highly clustered habitat, while Row 2 (Panels C and D) shows the performance of RSFs in a scenario of unclustered habitat. Black points in Column 1 (Panels A and C) indicate mean parameter estimates for weighted RSFs performed on 400 simulated movement tracks, while white points in Column 2 (Panels B and D) indicate mean parameter estimates for IID RSFs. Bars indicate mean confidence intervals for selection of Habitat 1 over Habitat 2 across all 400 simulated movement tracks. In this case, the animal is not selecting habitat, yet 50% habitat-specific triangulation failure creates the appearance of avoidance of Habitat 2 in favor of Habitat 1. In both habitat scenarios, confidence intervals become narrower as sampling rate increases for IID RSFs, while weighted RSF confidence intervals remain stable. Parameter estimates improve (i.e., become closer to the truth at zero) for wRSFs as autocorrelation of the data increases, while parameter estimates of IID RSFs remain stable. The advantages of weighted RSFs are more clearly visible in the scenario of highly clustered habitat, but weighting still improves both parameter estimates and confidence intervals when habitat is not strongly clustered.

### 3.2 Empirical Examples

When analyzing the water mongoose data (Fig. 3A), we found that both IID and weighted RSFs identified statistically significant avoidance of plantation areas, but the point estimate of avoidance of plantation areas for the weighted RSF lies closer to zero and outside the 95% confidence interval generated by the IID RSF (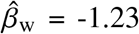 vs. 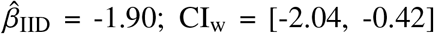 vs. CI_IID_ = [-2.35, -1.45]). Furthermore, although point estimates for avoidance of built-up areas were similar between weighted and IID RSFs (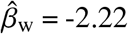 vs. 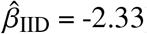), avoidance of built-up areas was only statistically significant for the IID RSF (CI_w_ = [-4.79, 0.35] vs. CI_IID_ = [-3.66, -1.01]).

When analyzing the caracal data (Fig. 3B), we found that although the IID RSF identified statistically significant avoidance of plantation areas 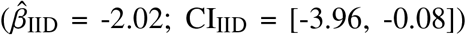, the weighted RSF found little evidence for avoidance 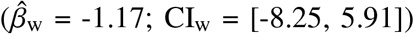. Moreover, although point estimates for selection of built-up areas were similar (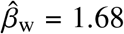 vs. 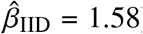), confidence intervals were substantially wider for the weighted RSF (CI_w_ = [0.67, 2.69] vs. CI_IID_ = [1.39, 1.78]).

When analyzing the serval data (Fig. 3C), we found that although point estimates for avoidance of plantation areas were similar between weighted and IID RSFs (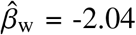 vs. 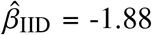), avoidance of plantation areas was only statistically significant for the IID RSF (CI_w_ = [-5.65, 1.56] vs. CI_IID_ = [-2.47, -1.30]). Point estimates for avoidance of built-up areas were also similar between weighted and IID RSFs (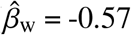 vs. 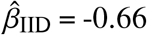), but this avoidance was again only statistically significant for the IID RSF (CI_w_ = [-1.88, 0.73] vs. CI_IID_ = [-0.90, -0.43]).

When analyzing how data thinning would influence inferences of habitat selection by the caracal, we found that IID RSFs parameterized using subsets of the caracal movement track exhibited substantial variation around the parameter estimate provided by the weighted RSF (Fig. 5). The parameter identified by the weighted RSF fell within 27 out of 27 (100%) of the IID RSF 95% confidence intervals, but 7 out of 27 (26%) of the IID RSFs did not identify statistically significant habitat selection for this individual.

**Figure 4:**
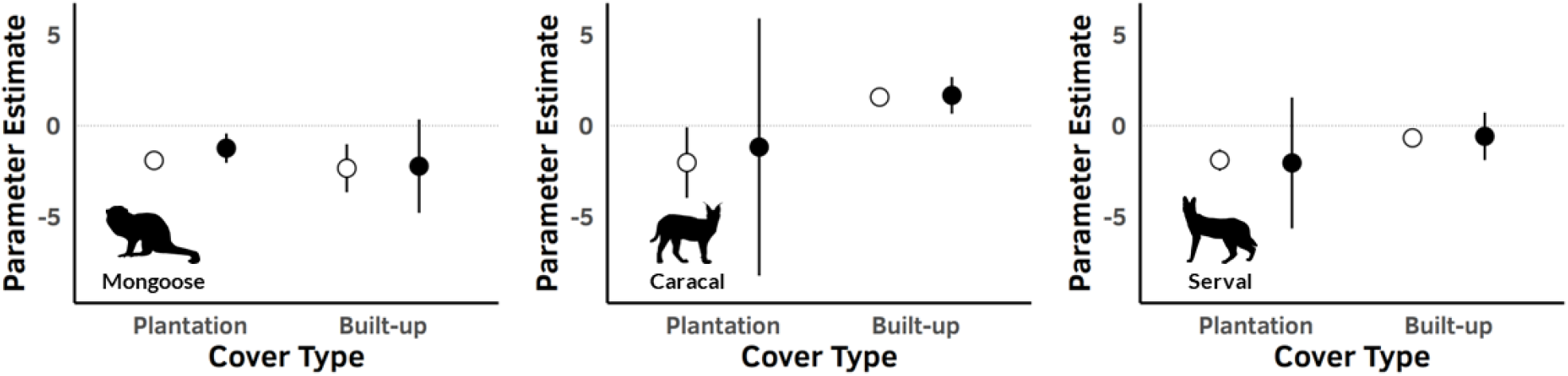
Parameter estimates and confidence intervals for IID (white points) and weighted (black points) RSFs during empirical tests on a water mongoose (Panel A), a caracal (Panel B), and a serval (Panel C). Weighting meaningfully alters point estimates of selection for built-up areas for the mongoose, and appropriately expands confidence intervals around parameter estimates, changing inferred habitat selection for all three animals.

**Figure 5:**
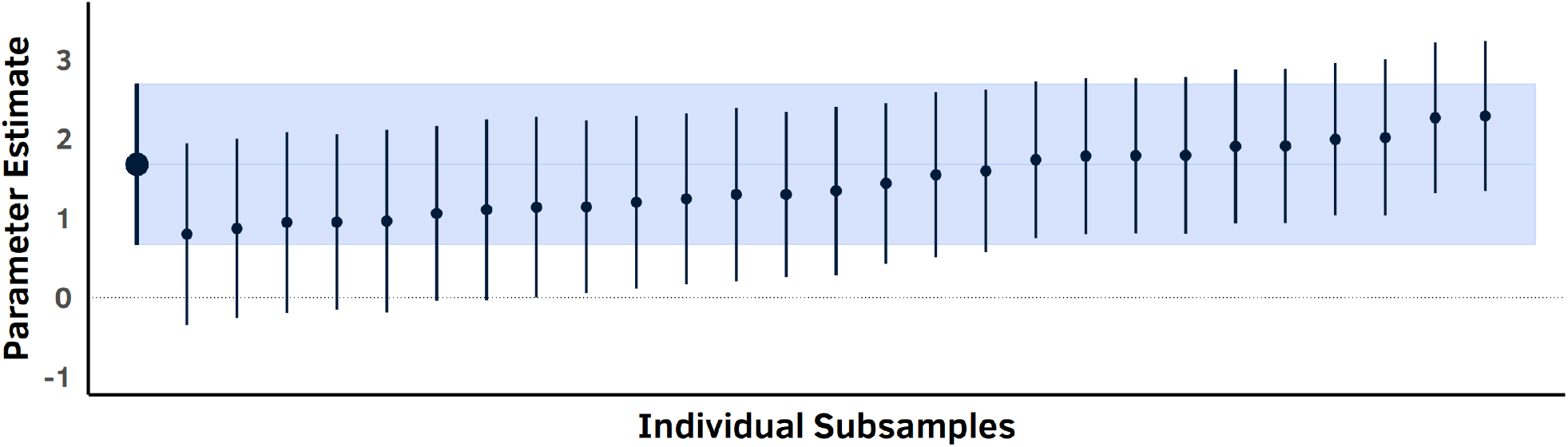
Parameter estimates and confidence intervals around those parameter estimates for one weighted RSF performed on a complete caracal movement track and 27 IID RSFs performed on subsamples of the movement track in which each sequential location was independent. The larger point and bar on the far left represent the weighted RSF, while the smaller points and bars represent individual subsets. The light blue box outlines the confidence interval of the weighted RSF, while the darker blue horizontal line identifies the selection parameter identified by the weighted RSF. As expected, IID RSFs parameterized using thinned data sets exhibit substantial variation in their estimated parameters.

Our empirical analyses demonstrate that inferences gained from weighted and IID RSFs can diverge substantially. Weighted RSFs produce wider confidence intervals than conventional RSFs and meaningfully different parameter estimates when missing data is common, indicating that the results of our simulations translate to real-world data, even at absolute sample sizes that are relatively small for modern GPS data sets. When data are autocorrelated, thinning data allows IID RSFs to generate more robust confidence intervals, but with more variation in parameter estimates than weighted RSFs.

## 4 Discussion

Although the autocorrelated nature of modern animal tracking data is known to pose statistical challenges for conventional RSFs that assume IID data (Boyce, 2006, Fieberg *et al*., 2010, Northrup *et al*., 2013), no generally accepted solutions to these challenges have been described. Wildlife biologists therefore often fit IID RSF models to autocorrelated tracking data, which can lead to both pseudoreplication and bias in parameter estimates of RSFs. In this study, we introduced a method of weighting individual locations to account for autocorrelation, and which reduces pseudoreplication and bias in RSFs. This method can improve estimates of confidence intervals provided by RSFs when tracking data are autocorrelated (as is the case for most modern animal tracking data sets), dramatically reducing the incidence of Type I error in studies of resource selection. It also reduces bias in parameter estimates when some covariates are over- or undersampled, such as when study animals inhabit areas where landscape features cause frequent triangulation failures in GPS devices, researchers employ duty cycling, or researchers use tracking technologies that do not produce regularly sampled location data (e.g., VHF telemetry or visual searches for marked individuals). In the future, our approach could be extended further to address other statistical problems associated with RSFs, which we discuss below.

Rigorous estimation of confidence intervals on RSF parameters is a challenge using conventional RSF methods. Because consecutive data points in modern animal tracking data are rarely statistically independent, using all the data available in an autocorrelated movement track leads to increasing (and increasingly misleading) confidence in parameter estimates with increasing sampling rates. Data thinning can be used to reduce autocorrelation between successive locations, which can widen confidence intervals that are known to be too narrow. In practice, however, data thinning inherently involves loss of useful information on an animal’s movements, is generally applied in an ad-hoc and non-optimized way, and can pose other statistical problems (Noonan *et al*., 2020). Generalized estimating equations can also be used to estimate more robust confidence intervals while using all the data in a movement track, but the “cluster” sizes (akin to effective sample sizes) are often also chosen arbitrarily (Prima *et al*., 2017), and it is difficult to implement generalized estimating equations jointly with other common techniques to mitigate statistical issues in RSFs (e.g., random effects). The method we propose in this study offers an objective way to estimate effective sample sizes for generating reliable confidence intervals.

In addition to estimating more robust confidence intervals, our method of weighting also offers potential to mitigate bias that can arise from inadequate sampling of landscape features that cause triangulation failure, irregular sampling designs intended to document animal behavior at multiple temporal scales, or technologies that do not allow regular sampling intervals. As shown by our simulations (Table 1; Fig. 3), RSFs that assume IID data cannot handle frequent habitat-induced triangulation failure—estimates of selection of those habitats are negatively biased, and models become artificially more confident in this bias as the sampling rate increases. Our method of weighting reduces this bias, and the quality of estimated parameters improves rather than declines as the sampling rate increases (however, we note that the extent to which inference is improved is dependent upon the spatial arrangement of environmental covariates). Duty cycling, by which researchers periodically suspend or reduce the sampling rate in an attempt to collect high-resolution data over longer total periods of time while conserving battery life, offers another promising use case for our method. For example, researchers may be interested in the foraging behavior of a crepuscular animal and therefore program a tracking device to collect more locations near dawn and dusk, when an animal is actively foraging. In this case, an IID RSF would result in sampling foraging areas out of proportion to other areas where an animal spends its time, potentially underestimating use of landscape features that play important roles as rest sites, escape cover, and/or thermal refugia. A weighted RSF could down-weight oversampled crepuscular locations more than undersampled locations during the day and night, providing more objective parameter estimates for overall resource selection. Finally, many species are not amenable to wearing tracking devices that acquire fixes at regular intervals. Aquatic or semi-aquatic species may spend long periods of time underwater where they are inaccessible to satellites (Breed *et al*., 2011, Fleming *et al*., 2018), and many species remain too small for today’s GPS technology (McMahon *et al*., 2017, Weller *et al*., 2016). Weighted RSFs provide a way for researchers to conduct RSFs that are less biased by such technological constraints.

Although applied here to tracking data of individual animals, the statistical principles underlying our method are also applicable at the species and population levels, which would be of great use to researchers. Random effects are often used to account for non-independence of data for individuals within a population. Gillies *et al*. (2006) recommended using random intercepts to account for unequal sample sizes between individuals in studies of habitat selection, and random slopes to account for differences in habitat selection between individuals. Similarly, Hebblewhite & Merrill (2008) recommended using random intercepts to account for non-independence arising from correlated locations in socially structured populations (e.g., repeated observations from individuals within the same herd or pack). Incorporating random effects into RSFs and SSFs has since become standard practice (Fieberg *et al*., 2010, Muff *et al*., 2020), but remains an imperfect solution for accounting for pseudoreplication. Random effects generate confidence intervals that depend on variation in the point estimates of RSF parameters between individuals, which may or may not reflect true parameter uncertainty. If autocorrelation in individual movement tracks is not properly accounted for, individual sampling variance is underestimated, which inflates population variance estimated using random effects. Checking for this is difficult and rarely (if ever) performed. Population-level likelihood weighting, however, has potential to serve similar purposes. Weighting based on effective sample sizes can account for individuals having been tracked for differing amounts of time, and autocorrelation-informed weighting can account for differing amounts of spatial or temporal autocorrelation between individuals in a study area or study period. Iterated over both individuals and populations, such weight optimization accounts for temporal sampling bias within individual time-series, as it did for the single-individual weighting described in this study, while also accounting for sampling biases among individuals, where certain individuals are better sampled than others.

Because location weights are generated from range distributions using weighted Autocorrelated Kernel Density Estimation, which can only be validly calculated on animals that are range-resident, this method cannot be used to conduct RSFs on individuals that are not range-resident. However, it should be noted that because conventional IID RSFs assume stationary distributions of resource use and availability, conventional RSFs also assume range residency (but this assumption is often ignored). Step-selection functions are therefore more appropriate for studying resource selection by migratory, dispersing, or nomadic animals because they do not assume stationary distributions of use and availability. The inferences gained from such studies should be treated with caution, however, because autocorrelation can still lead to pseudoreplication and bias in inferences gained from SSFs of these movement tracks.

Although ecologists and conservation biologists have access to better statistical and computational tools than ever before, new methods are still required to maximize the value of animal tracking data for informing our understanding of the natural world. In this study, we introduced a method of autocorrelation-informed weighting to reduce pseudoreplication and bias when parameterizing RSFs with autocorrelated and irregular animal tracking data, which are pervasive in studies of animal movement. Our method is easily implemented using the `ctmm` R package (Calabrese *et al*., 2016), and we provide an annotated workflow in our supplementary materials (Appendix 1) so that other researchers can implement weighted RSFs on their own animal location data. We hope that the analytical technique we have provided here, which is grounded in statistical theory and validated using simulations and empirical data, allows continued progress toward more statistical rigor in studies of resource selection by animals.

## Acknowledgements

We thank I. Silva for providing feedback on the weighted RSF tutorial in Appendix 1. JC and CF were supported by NSF IIBR 1915347. RK was supported by NSF ABI 1914928 and NASA Ecological Informatics 80NSSC21K1182. This work was partly funded by the Center for Advanced Systems Understanding (CASUS) which is financed by Germany’s Federal Ministry of Education and Research (BMBF) and by the Saxon Ministry for Science, Culture and Tourism (SMWK) with tax funds on the basis of the budget approved by the Saxon State Parliament.

## Data Accessibility

All analytical code related to this manuscript is available at https://github.com/integratecology/wrsf. All data and code used in analyses for this manuscript will be made publicly available on the *RODARE* data repository upon publication of this manuscript.

## Authors’ Contributions

JC and CF conceived the ideas; CF led the mathematical derivation and software implementation; JA conducted the simulations and empirical analyses; JS, CD, and TR contributed empirical data; and JA and CF led the writing of the manuscript. All authors contributed critically to drafts of the manuscript and gave final approval for publication.

## Appendix S2

**Figure S1:**
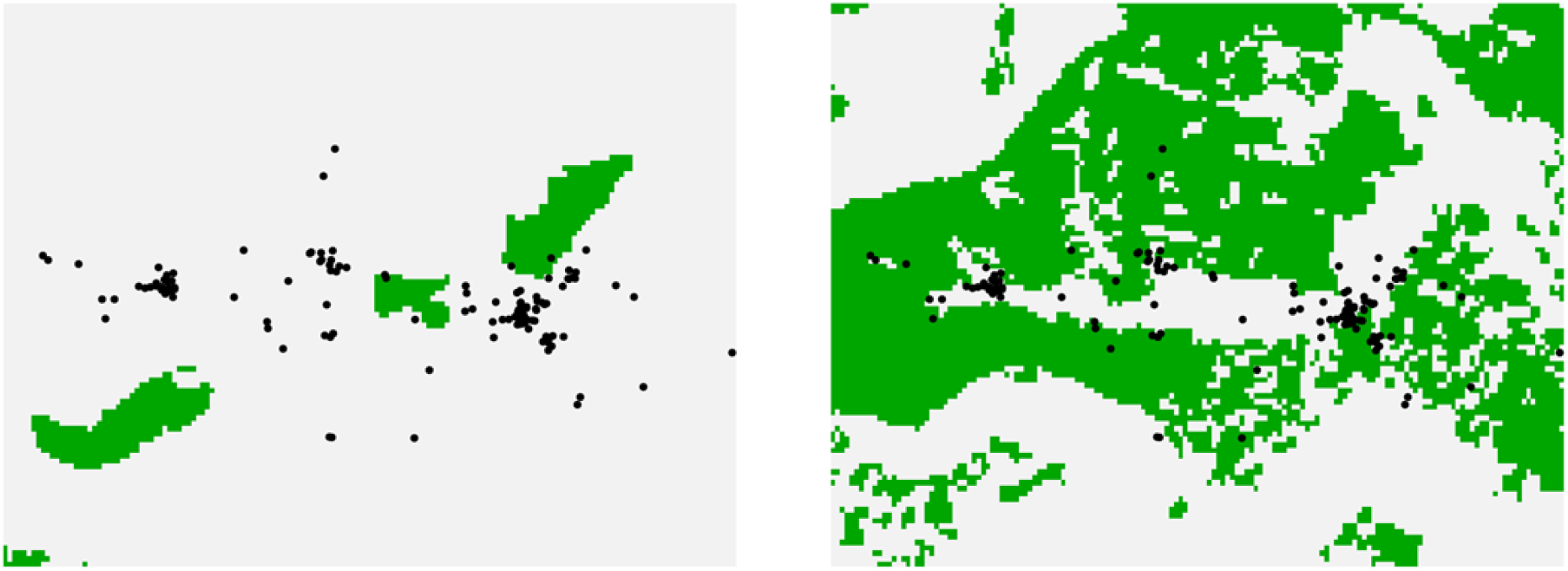
The movement track for the water mongoose used in our empirical example, super-imposed on the plantation (left) and built-up (right) cover types mapped by EKZN Wildlife & GeoTerraImage (2018). Green represents the cover type of interest.

**Figure S2:**
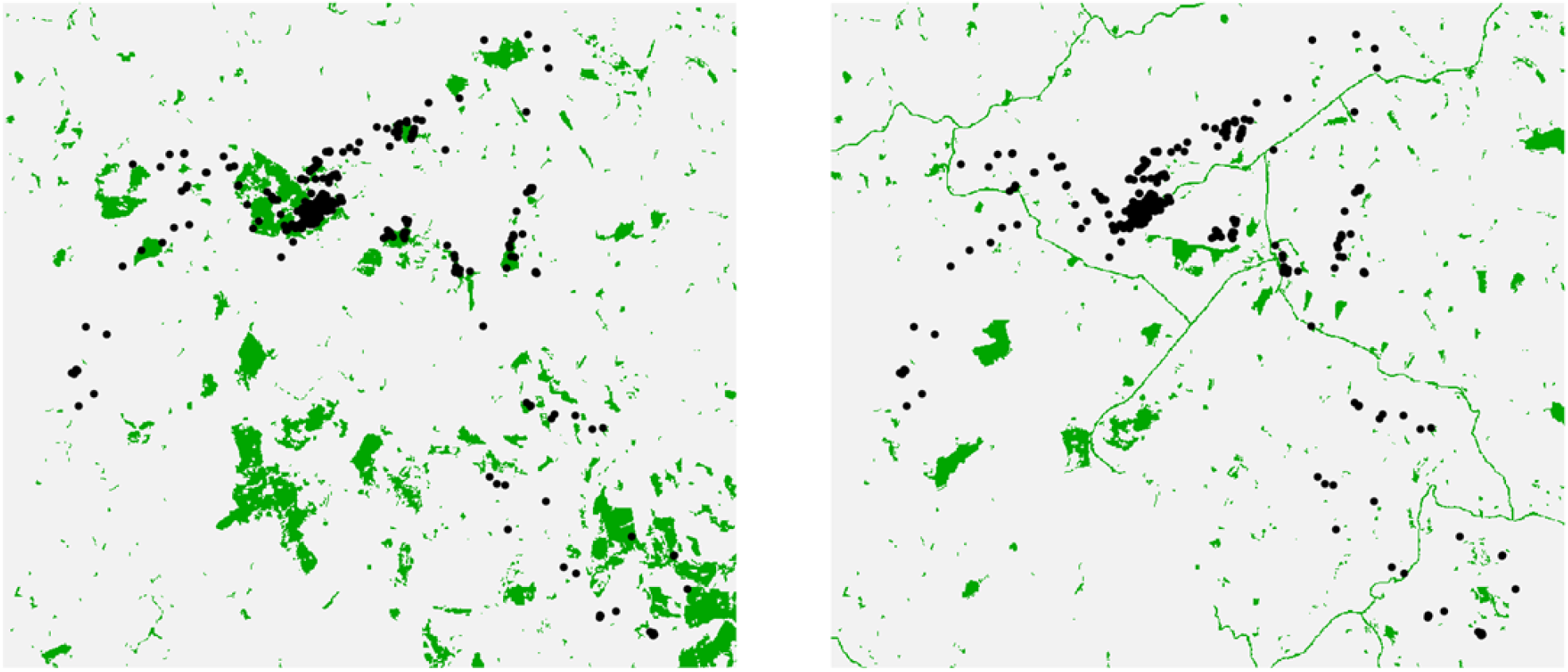
The movement track for the caracal used in our empirical example, superimposed on the plantation (left) and built-up (right) cover types mapped by EKZN Wildlife & GeoTerraImage (2018). Green represents the cover type of interest.

**Figure S3:**
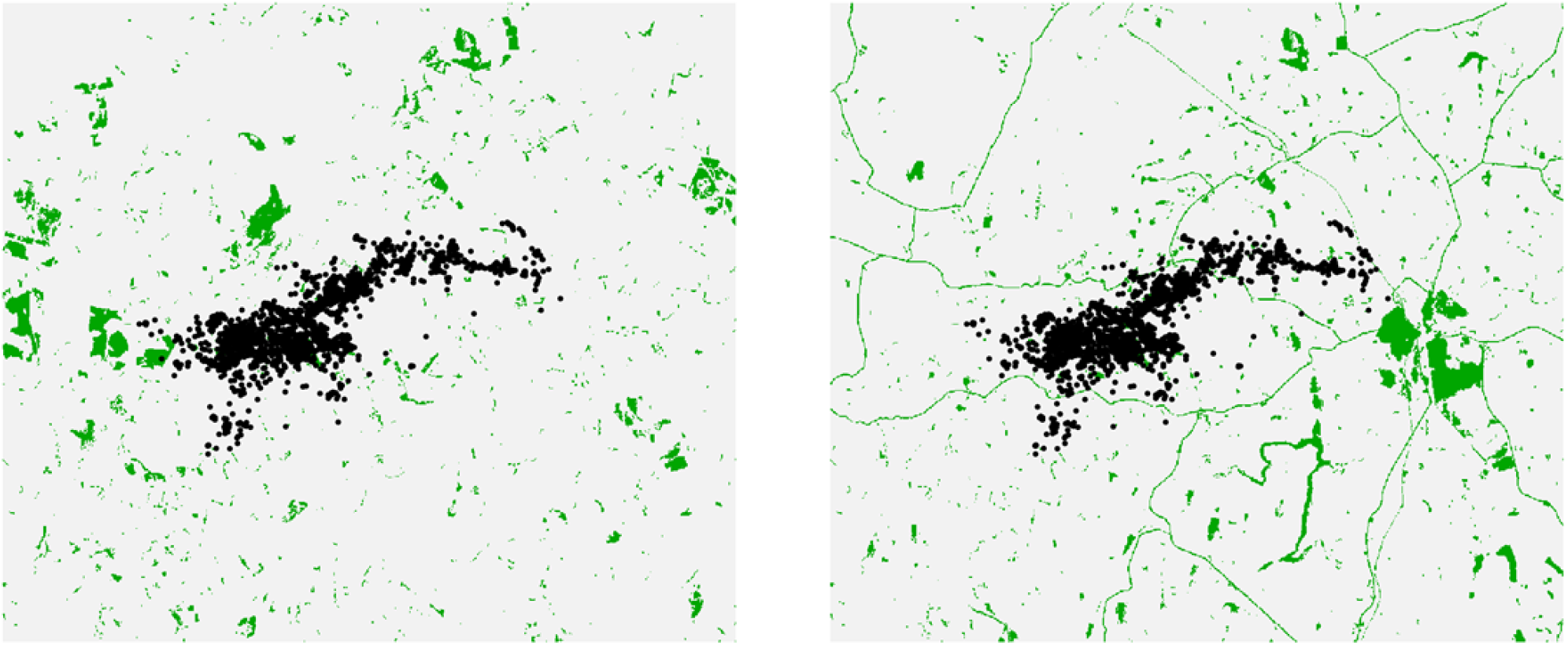
The movement track for the serval used in our empirical example, superimposed on the plantation (left) and built-up (right) cover types mapped by EKZN Wildlife & GeoTerraImage (2018). Green represents the cover type of interest.

## Appendix S3

### Availability Sampling with Integrated RSFs

This appendix describes how we sampled availability for our simulations and empirical analyses in this paper. The following text is excerpted from the current draft of an in-preparation manuscript (Fleming et al., in prep) on the topic of availability sampling in RSFs.

### Conventional RSF Analysis

Although RSFs are typically fit using logistic regression, biometricians usually model RSFs as an inhomogeneous Poisson point process (Aarts *et al*., 2012, Fieberg *et al*., 2021, Fithian & Hastie, 2013, Warton & Shepherd, 2010). Such inhomogeneous Poisson point process models allow sampling of availability to be mathematically derived by numerically estimating a normalization integral of the model likelihood (Warton & Shepherd, 2010), which leads to parameter convergence with sufficiently large quantities of available locations (Fieberg *et al*., 2021). Below, we outline the mathematical definitions that characterize this approach, and then our “integrated” method.

#### Inhomogeneous Poisson Point Process Models

For a single individual tracked along a movement path, the marginal density of a location (*x, y*) sampled from an inhomogeneous Poisson point process (iPPP) with intensity function *λ*(*x, y*) can be represented by:

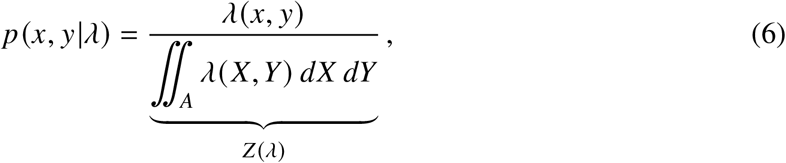

where *A* is the *domain of integration* and *Z* (*λ*) is the normalization constant. For the Poisson point process, the intensity function *λ*(*x, y*) determines the rate of events (e.g., animal visits) at a location given the history of events in a stationary process. Equation (6) is the basic RSF model for typical GPS or VHF telemetry data: time-series data for a fixed count of one, which has a random location. The exponential model is usually used to describe resource selection, so under a log link function the intensity function can be represented by:

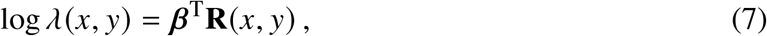

where **R**(*x, y*) represents spatially explicit covariates or resources, and ***β*** are their corresponding regression parameters. The probability density function (6) can therefore be represented as:

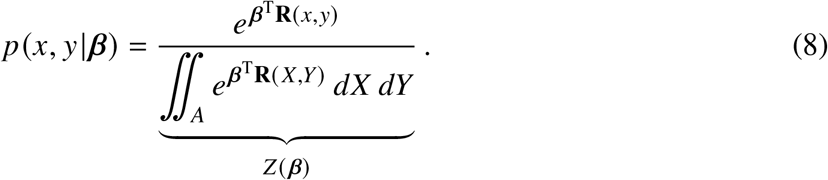

#### Proper Distributions

Importantly, (8) must be a *proper distribution*, meaning that the denominator, *Z* (***β***), must evaluate to a finite normalization constant that fixes the total probability mass to 1. Traditionally, this has been imposed by choosing a finite domain of integration, *A* (typically interpreted in biological terms as the area available to an animal; hereafter, “available area”). The conventional model is therefore given by

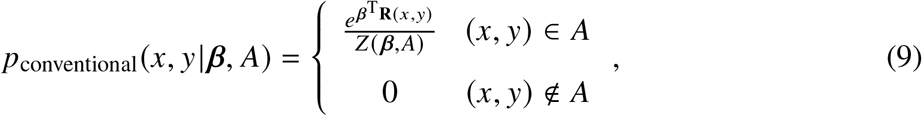

which is explicitly a function of the available area *A*. This means that any animal locations outside of *A*—which may be many locations if *A* includes less than 100% of the area an animal travels—necessarily occur in areas where the probability of use is assumed to be zero.

#### Monte-Carlo Integration

The normalization integral *Z* (*λ*) in (1)—or *Z* (***β***) in (3)—can be approximated numerically using Monte Carlo integration, which makes use of random sampling within a known parameter space to numerically estimate integrals of interest. In RSFs, this occurs by sampling locations within the available area (hereafter, “availability sampling” of “available locations”). In terms of available locations in the available area,

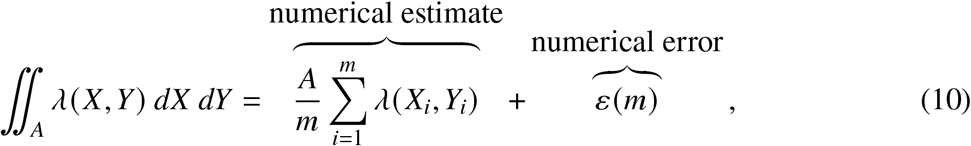

where *m* denotes the number of available locations, (*X*_*i*_, *Y*_*i*_), that are randomly sampled from the uniform distribution with area *A*. The variance of the numerical error decreases with the number of available locations, *m*, but increases with the available area, *A*, given the relation

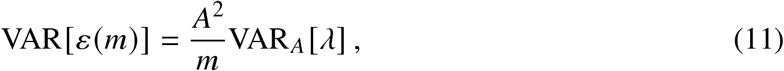

in terms of the spatial variance of the intensity function, VAR_*A*_ [*λ*], which can be estimated from the available locations according to

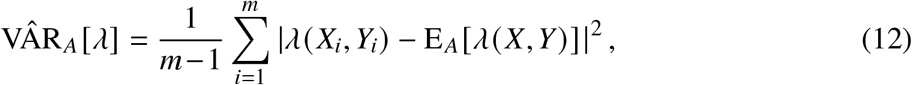

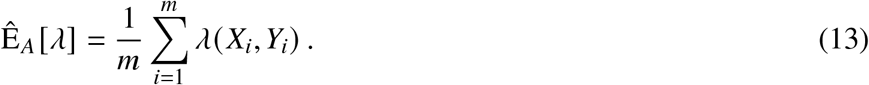

With some rescaling, the likelihood function of the data can be represented as that of a conditional logistic regression under a logit-link, which is the prevalent strategy for parameterizing SSFs and iSSAs (Avgar *et al*., 2016, Fortin *et al*., 2005, Signer *et al*., 2019). In the limit of infinite *m*, the logistic regression parameter estimates converge to those of the iPPP model (Aarts *et al*., 2012, Fithian & Hastie, 2013, Warton & Shepherd, 2010).

### Integrated RSF Analysis

In contrast to the conventional two-stage method for parameterizing RSFs, it is also possible to include spatial terms in the RSF model that determine the domain of availability simultaneously with model fitting. Although the value of explicitly incorporating area in RSF models has been recognized before (Fieberg *et al*., 2021), the benefits of this approach are not widely appreciated among ecologists who conduct resource selection studies, and in practice, spatial terms are rarely, if ever, included in RSFs. Incorporating spatial terms into RSFs can also account for animal movement in the RSF model (albeit at coarser scales than iSSAs). Below, we describe our approach.

#### Integration of Availability Parameters

In an integrated RSF, we start with the same iPPP modeling framework (6), but rather than fixing availability as a uniform distribution within an arbitrary spatial boundary—as in (4)—we substitute a more gradually decaying intensity function *λ*(*x, y*), such that the integral

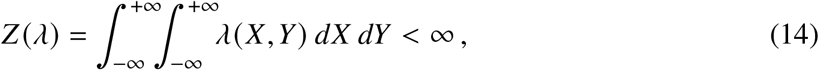

converges without ad hoc truncation. By explicitly modeling non-uniform availability with the intensity function *λ*(*x, y*), the availability parameters (which define the spatial extent of the area that is available to an animal) are estimated consistently and simultaneously with the resource-selection parameters. A number of different models can be used to achieve this goal, but we will use a symmetric Gaussian availability model by including three additional spatial covariates—*x, y*, and *x*^2^ + *y*^2^—or equivalently

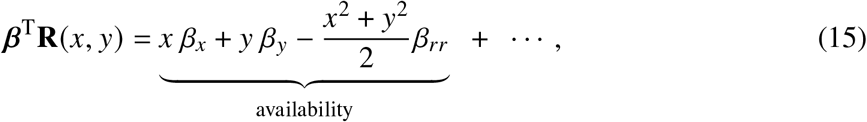

plus conventional resource-selection terms. In the absence of resource selection, this results in a Gaussian utilization distribution, with a mean location of E[(*x, y*)] = (*β*_*x*_, *β*_*y*_)/*β*_*rr*_ and variance of VAR[*x*] = VAR[*y*] = 1/*β*_*rr*_. This means that when an animal exhibits no selection for any covariates included in the model, the probability density of its location at any given time is still estimated as a symmetrical Gaussian utilization distribution, which serves as the null model. Availability parameters are thus estimated consistently with the resource-selection parameters, which increases the statistical efficiency of the fitted model, reducing bias in the parameter estimates and improving the coverage of their confidence intervals. In particular, because we do not treat *β*_*x*_, *β*_*y*_, and *β*_*rr*_ as known parameters on which to condition a second-stage of analysis, this approach also accounts for uncertainty in the unknown available area, by accounting for the correlation between errors in the availability-parameter estimates and resource-selection parameter estimates, and enabling estimation of confidence intervals around *A*.

## Notes

### Competing Interest Statement

The authors have declared no competing interest.

### Summary of Updates

This preprint has been revised to reflect alterations to the text throughout the manuscript and add additional simulations and empirical examples.

