## Appendix 1 for "Mitigating pseudoreplication and bias in resource selection functions with autocorrelation-informed weighting": appendix1.Rmd

````
---
title: "Appendix 1: Weighted resource selection functions in the 'ctmm' package"
author: "Jesse M. Alston*, Christen H. Fleming*, Roland Kays, Jarryd P. Streicher, Colleen T. Downs, Tharmalingam Ramesh, and Justin M. Calabrese"
date: "`r Sys.Date()`"
output: rmarkdown::html_vignette
---

In this R tutorial, we will take you through data preparation and fitting of weighted resource selection functions using the `ctmm` package.

# Step 1: Data Preparation

In this tutorial, we will use data from an African buffalo (Syncerus caffer) that is already available in the `ctmm` package. Our example data is therefore already prepared as a list of `telemetry` objects. If you want to import raw location data but have not done this before, you can find detailed information about how to do this at: https://ctmm-initiative.github.io/ctmm/articles/variogram.html. The code chunk below loads and plots location data from the buffalo we will use.

```{r,  fig.show = 'hold', message = FALSE}
# At the moment, weighted RSFs require the development version of ctmm, 
# which can be installed using the "remotes" package
# remotes::install_github("ctmm-initiative/ctmm")

# Load the ctmm package
library(ctmm)

# Load buffalo data
data(buffalo)

# Select an individual buffalo
pepper <- buffalo$Pepper

# Plot the data
plot(pepper)
title("Buffalo GPS Locations")
```

# Step 2: Fitting a Continuous-time Movement Model

Selecting and fitting a continuous-time movement model is the first step in almost every analysis in the 'ctmm' package, and weighted resource selection functions are no different. Detailed information on how to do this is available at: https://ctmm-initiative.github.io/ctmm/articles/variogram.html. (Warning: In this example, this will take a few minutes).

```{r,  fig.show='hold'}
# Calculate the variogram
SVF <- variogram(pepper)

# Estimate an initial model
GUESS <- ctmm.guess(pepper, variogram=SVF, interactive=FALSE)

# Select the best model
FIT <- ctmm.select(pepper, GUESS, trace=2)

# Examine the model outputs
summary(FIT)
zoom(SVF, FIT)
```

# Step 3: Calculating location weights

The next step is calculating the likelihood weights for each location. These weights are a good measure of how autocorrelated each data point is, and are borrowed from a weighted AKDE home range estimate.

```{r,  fig.show='hold'}
# Calculate a weighted AKDE home range
ud <- akde(pepper, FIT, weights=TRUE)

# Examine the outputs
plot(pepper, ud)
summary(ud)
```

# Fitting a weighted resource selection function

The last step is fitting a resource selection function to a habitat raster. We will create an arbitrary east-to-west gradient for this example, but you can import any raster data set using the raster() function in the 'raster' package. In this example, the raster is generated independently of the movement track, so we find that there is no selection.

```{r,  fig.show='hold'}
# Load raster package
library(raster)

# Create a habitat raster
r1 <- raster(nrows = 100, ncols = 100, 
             xmn = 31, xmx = 32.5, ymn = -25, ymx = -23.5,
             vals = c(rep(seq(1, 100,1),100)))
projection(r1) <- "+proj=longlat +datum=WGS84 +no_defs"

# Plot raster to see where it is in relation to the buffalo
raster::plot(r1)
points(pepper$latitude~pepper$longitude)

# Weighted resource selection function with one habitat covariate
wRSF <- ctmm:::rsf.fit(pepper, UD=ud, R=list(habitat1=r1), error=0.1)
summary(wRSF)

```

The four arguments in the rsf.fit() command are the movement track (a telemetry object), the utilization distribution (inputting a weighted AKDE automatically makes the RSF a weighted RSF), a list of habitat rasters, and the amount of numerical error you decide is acceptable (more details on numerical error in RSF parameterization are forthcoming in a future paper).

The outputs that will be most valuable to most users are the effective sample size (which can be located using summary(wRSF)$DOF["area"]) and the model coefficients (in this case, the top row of the $CI table, which can be located using summary(wRSF)$CI[1,]).
````
